# Multimodal joint deconvolution and integrative signature selection in proteomics

**DOI:** 10.1101/2023.10.04.560979

**Authors:** Yue Pan, Xusheng Wang, Chunyu Liu, Junmin Peng, Qian Li

## Abstract

Deconvolution is an efficient approach for detecting cell-type-specific (cs) transcriptomic signals without cellular segmentation. However, this type of methods have not been extended to the proteomics research. Here we present a novel algorithm and tool to dissect bulk proteome by leveraging the information shared between transcriptome-proteome. Our tool first identifies potential cell marker proteins by integrating RNA and protein bulk expression profiles and then jointly quantifies the cell abundance in mixture proteomes without using a reference signature matrix, enabling the downstream analyses such as cs-protein Quantitative Trait Loci (cspQTL) mapping. This new method and the cspQTL analysis are implemented in the R package MICSQTL that also provides integrative visualization of bulk multimodal samples, available at https://bioconductor.org/packages/MICSQTL.

## Introduction

Proteomics profiling and analysis at cell type level is critical in the study of complex biological systems with numerous applications in immunology, cancer research, and developmental biology [1–3]. Several technologies have been developed to identify and quantify proteins at cellular resolution [4]. For example, the detection of proteins in single-cells by CITE-seq [5], which is a multimodal sequencing technique that enables simultaneous profiling of gene expression and up to 300 cell surface protein markers in individual cells, allows the identification of rare cell types and cells that express low levels of certain genes. However, this technology only measures limited number of proteins.

Recent advances in liquid chromatography mass spectrometry (LC–MS)-based proteomics methods have addressed the limitations in the sensitivity and throughput [6, 7], which accelerates the evolvement of single cell mass spectrometry (scMS) proteomics. One major challenge in scMS proteomics is that the number of unique samples and cells analyzed in a single day is very limited [8]. For label-free scMS, samples are analyzed sequentially with analysis time ranging from 35 to 90 min. At this speed, a maximum of 40 single cells could be analyzed in a day, which is not ideal for population-scale clinical studies due to the burden of time and costs. The scMS technology limitations and costs of cell sorting are the hurdles for cell-type-specific inference such as differential expression (csDE) or protein quantitative trait mapping (cspQTL) that requires median or large sample size.

Deconvolution algorithms are rapidly developed to measure molecule proportions (e.g., RNA transcripts) mapped to each cell type, which is different from the cell counts composition and varies across molecular sources. To estimate the cellular composition in human proteome, the pure cell or single cell reference proteomes (i.e., signature matrix) is needed but lacking due to the challenges in cellular dissociation for certain tissue or cell types (e.g., astrocytes and exitatory neurons [2]) and the aforementioned limitations in scMS and CITE-seq. Meanwhile, the multi-omics profiling matched by samples becomes popular in recent decade, enabling the integration across data sources and the discovery of multimodal signatures for disease [1]. Hence, we design a novel algorithm to estimate the proteomics cell fractions by integrating bulk transcriptome-proteome without reference proteome, implemented in R package MICSQTL. Our method enables the downstream cell-type-specific protein quantitative trait loci mapping (cspQTL) based on the mixed-cell proteomes and pre-estimated proteomics cellular composition (Figure 1), without the need for large-scale single cell sequencing [9] or cell sorting.

**Fig. 1.**
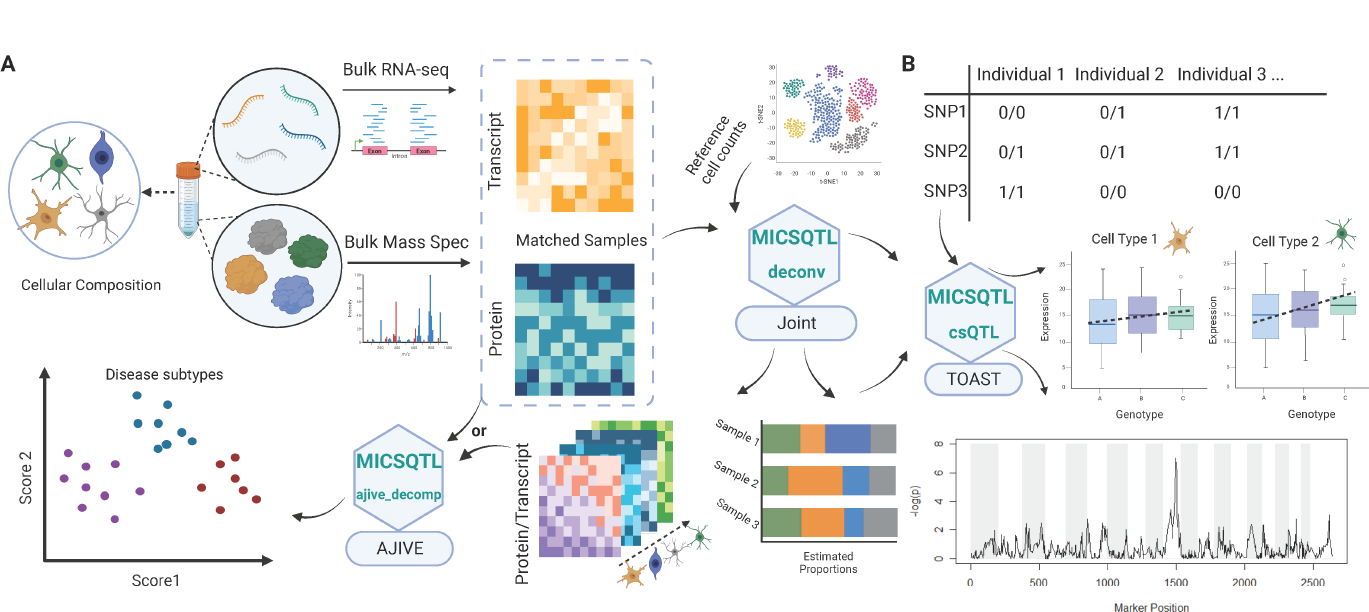
Flowchart of MICSQTL (A) Cross-source joint deconvolution and integration of matched transcriptome-proteome. (B) Cell-type-specific pQTL based on estimated cell fractions.

## Results

### MICSQTL algorithm and implementation

For each tissue sample with bulk transcriptome-proteome, we model the cellular compositions ***θ***^(*i*)^ in the *i*th modality as a product of tissue-specific cell counts fractions ***p*** and molecule source-specific cell size factors ***s***^(*i*)^. An unobserved individualized reference panel is incorporated in this framework to allow inter-subject heterogeneity per cell type. We employ a loss function that integrates the observed bulk RNA and protein expressions to optimize the cell abundances in each molecular source, as described in Methods. The parameters are estimated by Projected Gradient Descent (PGD), with initial values sampled from the bulk transcriptome-proteome and an external reference of cell counts in similar tissue type. The reference cell counts are usually accessible in small-scale single cell or flow cytometry experiments. Hence, this joint deconvolution algorithm is semi reference-free without using a signature matrix from scRNA-seq or pure cell profiling, implemented in the function *deconv*. An alternative approach to quantify proteome cell fractions is the algorithm in Tensor Composition Analysis (TCA) [10]. However, TCA requires the input of sample-wise pre-estimated RNA proportions.

In proteomics deconvolution, researchers may not have *a priori* knowledge about the cell marker proteins for certain tissue types, but the cell marker genes in transcriptome have been broadly identified and curated in public databases [11]. Here, we use the AJIVE framework [12] to construct a common space shared across two molecular sources: bulk RNA expression of cell marker genes and the sample-matched whole proteome, which captures the between-sample heterogeneity caused by cellular abundance variation. Next, the observed whole proteome is projected onto this shared space by employing the reduced-rank loadings from AJIVE, where protein dimensions and annotations are unchanged. The rank of loadings is determined by an inherent algorithm. The potential cell marker proteins are selected based on the feature-wise Euclidean distance between the projected and observed proteomes. This cross-source feature selection is similar to ReFACTor [13], but ReFACTor is only applicable to single modality and sets the rank of loadings by the assumed number of cell types. Hence, we name our signature selection procedure as ‘AJ-RF’ and use the selected proteins to improve joint deconvolution, implemented in the function *ajive decomp*.

### Validation using multimodal expression from CITE-seq

We first validate our algorithm by using the pseudo bulk multimodal expression profiles built from a public CITE-seq dataset with 161,764 human peripheral blood mononuclear cells (PBMCs) collected from eight donors [1], processed by 10 *×* 3^*′*^ technology and Seurat v4. For each single cell, the expression of 228 surface proteins and more than 30,000 RNA genes were measured, whereas 44 genes were mapped to the surface proteins. Hence, we generated pseudo bulk expression data for the overlapped RNA features and 228 surface proteins by aggregating feature-wise Unique Molecular Identifier (UMI) counts across the cells per donor. The ground truth of cellular fractions can be achieved by the annotated cell labels, which is identical for RNA and protein in CITE-seq.

We selected four donors (samples P1, P4, P5, P7 in Figure 2 (A)) with disparities in B cell, natural killer (NK) cell, dendritic cell (DC) and other T cells abundances to reconstruct the pseudo reference multi-subject cell counts required in the joint deconvolution. The cell counts fractions quantified by our algorithm 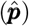 was similar to the true cell counts fractions captured and annotated in CITE-seq (r=0.8) as shown in Figure 2 (B), while the individualized cell-type-specific protein expression level resolved by our algorithm was significantly correlated with the observed pure cell bulk expression in three abundant cellular populations: CD4, CD8 T cells and monocytes (Figure 2(C)). This result demonstrated the validity and power of our algorithm, albeit the present work only focused on the cell abundance quantification in proteins.

**Fig. 2.**
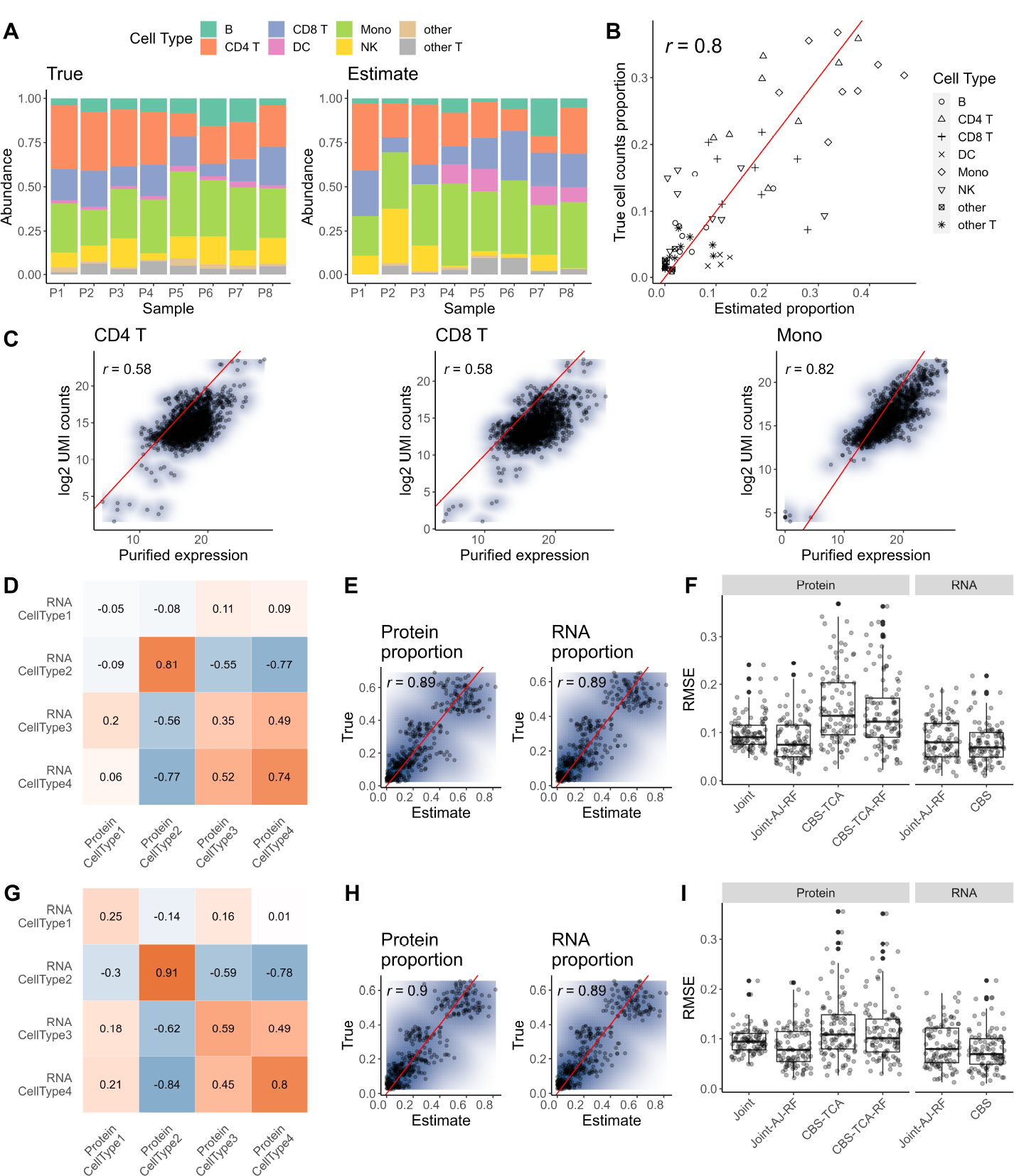
Performance of joint deconvolution in pseudo bulk or simulated multimodal expression. (A) Composition plot of true cell counts fractions (left) and estimated cell counts fractions (right). (B) Scatterplot comparing true cell counts fractions vs. jointly estimated cell counts fractions. (C) Scatterplots comparing the total UMI counts at log2 scale vs. purified expression level of each surface protein per sample in CD4 (left), CD8 (middle), and Monocytes (right). (D-F) Correlation coefficients between synthetic protein and transcript proportions per cell type (D); ground truth vs estimated cellular proportions per molecular source (E); Root Mean Square Error (RMSE) for different deconvolution methods, using accurate pseudo signature matrix in CBS, CBS-TCA, CBS-TCA-RF (F) in simulation scenario A. (G-I) Results in simulation scenario B.

### Assessment using simulation data

The above pseudo bulk data from CITE-seq only provides the ground truth for cell counts fractions instead of the cell proportions in each molecule source. To mimic the possible differences in RNA vs. protein cellular compositions and compare the performance of distinct methods, we designed a simulation study to generate sample-matched bulk transcriptome-proteome with ground truth of molecule-specific cell fractions. The real data parameters and statistical models used in simulation are described in Supplementary File. We designed two scenarios to allow relatively low (scenario A) and high (scenario B) correlations between the true cellular proportions in protein vs. RNA, which were moderately or strongly correlated (*r >* 0.3) in Cell Types 2-4 and weakly or not correlated (*r <* 0.3) in Cell Type 1, as shown in Figure 2 (D,G). The joint estimate of molecular source-specific proportions with AJ-RF protein selection (Joint-AJ-RF) are compared to the ground truth and visualized in Figure 2 (E). We further compared this result to the RNA proportions estimated by CIBERSORT (CBS) [14] and the protein proportions estimated by TCA two-step approach (CBS-TCA).

To apply CBS to the synthetic bulk RNA-seq data, we generated a pseudo signature matrix with distinct accuracies by adding small vs. large random noises to the true subject-specific purified transcriptomes, respectively. The cell marker genes are selected by gene-wise coefficients of variation calculated from the true reference panel, while 300 signature proteins were selected by AJ-RF for joint deconvolution based on these marker genes. Figure 2 (F) shows that the protein cellular composition output from our new method is more accurate than CBS-TCA and can be further improved by AJ-RF feature selection, while the ReFACTor algorithm applied to single modality does not improve the TCA-estimated protein proportions significantly. The CBS estimate using a relatively-accurate reference is slightly better than our jointly-deconvoluted cell fractions in either RNA or protein, but this outperformance is not profound. The bias of CBS RNA proportions or CBS-TCA protein proportions solved by an inaccurate signature matrix is inflated compared to the joint deconvolution results in RNA and protein (Figures S1, S3), while Joint-AJ-RF always yields accurate estimates across different scenarios (Figure 2 (D-I)) and PGD step size values (Figures S2, S4).

### Application to bulk transcriptome-proteome in bipolar disorder and schizophrenia

We used *n* = 264 tissue-matched transcriptome-proteome from postmortem human brain prefrontal cortex in a study of bipolar disorder (BP) and schizophrenia (SCZ) to demonstrate the performance of MICSQTL. The details of human brain tissues, MS proteomics and RNA-seq transcriptomics profiling are available in Supplementary File. The signature genes used in each deconvolution method were selected from previous findings [15]. To the best of our knowledge, the single cell or pure cell proteomics data profiled from human prefrontal cortex excitatory neurons are not publicly available. The protein proportions quantified by Joint-AJ-RF algorithm (Figure 3 (D)) showed disparities in astrocyte and oligodendrocyte abundances between (BP and/or SCZ) cases vs the young controls at similar age, i.e., *<* 70 years. This result is consistent with previous findings about glia cell abundance in SCZ [16, 17]. A recent snRNA-seq study [18] reporting cellular subpopulations in human brain prefrontal cortex for SCZ vs. healthy controls (Figure 3 (B)) also validated our estimates in Figure 3 (A). However, the CBS RNA deconvolution with signature matrix built from scRNA-seq data of healthy human prefrontal cortex [19] failed to recover the expected case-control disparities or mean abundances in multiple cell types (Figure 3 (C)). We also applied CBS to this bulk human prefrontal cortex proteomes with a reference panel of four cell types from major regions of mouse brain [15], which neither decomposed the neuron cells into excitatory and inhibitory subpopulations nor recovered the expected cell abundances (Figure 3 (D)).

**Fig. 3.**
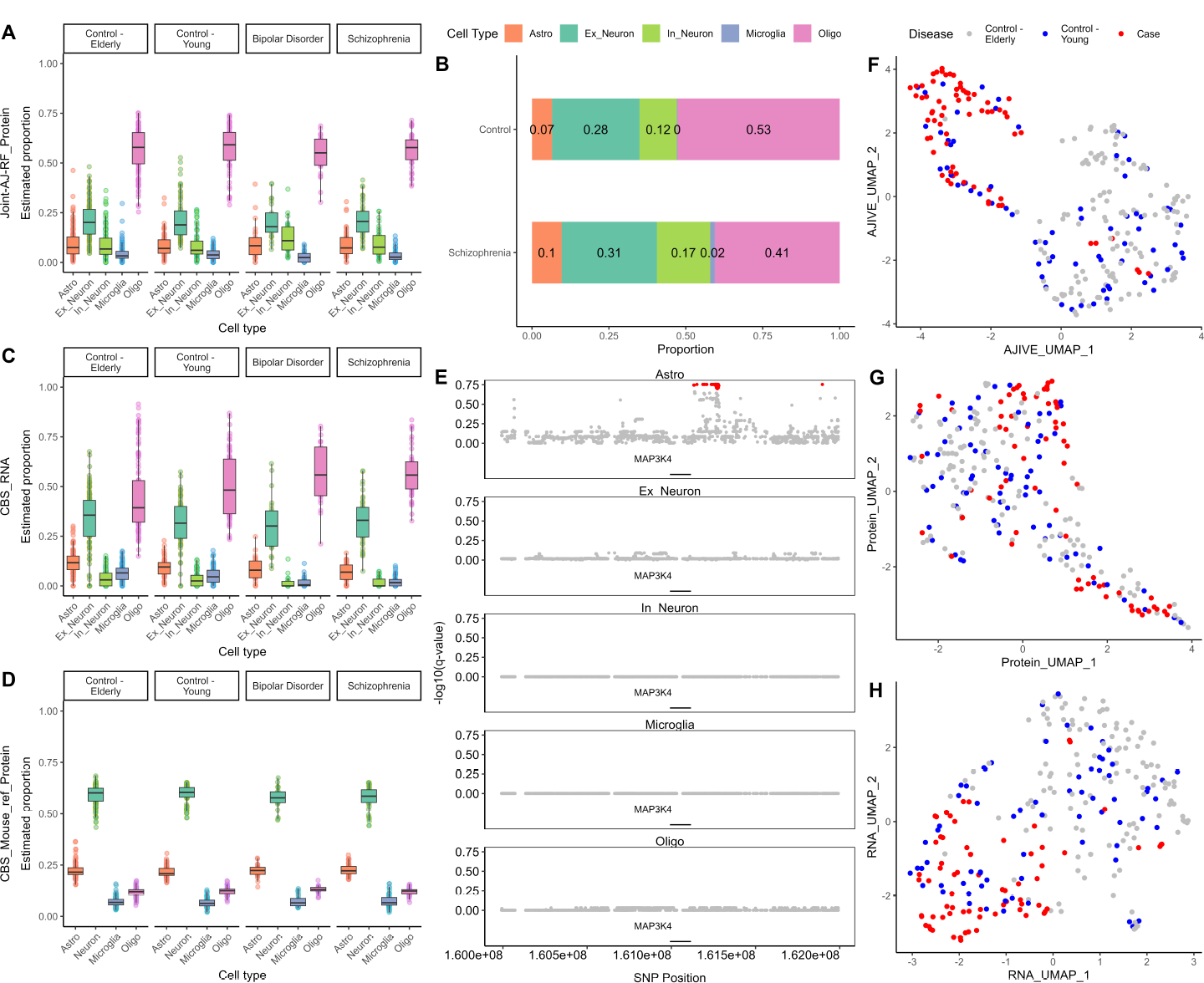
Performance of joint deconvolution with integrative signature selection. (A) Protein proportions in human prefrontal cortex quantified by Joint-AJ-RF. (B) Cellular composition measured by snRNA-seq for healthy controls and schizophrenia human prefrontal cortex. (C) Transcript proportions in human prefrontal cortex quantified by CIBERSORT with scRNA-seq reference in the same tissue type. (D) Protein proportions in human prefrontal cortex quantified by CIBERSORT with mouse brain pure cell reference. (E) FDR-controlled q-value for cspQTL analysis for protein *MAP* 3*K*4. (F)-(H) UMAP for joint and single-source visualizations.

A downstream cspQTL mapping was performed by cell-type-specific differential analysis, implemented in the function *csQTL*. The input data include bulk proteomes, whole genome sequencing genetic variants (SNP), and protein proportions in Figure 3 (A). For each protein of interest, we select the nearby SNPs within a 1D genomic distance range of 1 million bases (Mb). Figure 3 (E) shows the q-value with corrected false discovery rate per cell type for the protein *MAP3K4* coded by SCZ-associated genes [20]. More results of cspQTL analysis are visualized in Figure S4. Last but not least, our tool MICSQTL outputs the Common Normalized Scores (CNS) that represents the sample-specific variation shared across transcriptome and proteome with dimension determined by the common reduced rank of shared space. This multivariate scores may uncover the heterogeneity across disease phenotypes and the link to disease hidden drivers. The multi-omics human brain tissue samples is jointly visualized in Figure 3 (F) by concatenating the modalities with CNS, which outperforms the single modality visualization (Figure 3 (G-H)).

## Discussion

Our pipeline offers three primary functions to perform multi-omics cell abundance quantification with or without marker protein selection, integrative visualization, and cspQTL mapping. The semi reference-free joint quantification of cellular compositions in RNA and proteins were benchmarked by multiple datasets. That is pseudo bulk RNA and protein expression constructed from CITE-seq single cell data, synthetic bulk multimodal expression generated by statistical models, and real human brain bulk transcriptome-proteome with external snRNA-seq data.

Obviously, our joint deconvolution algorithm coupled with cross-modal signature protein selection yields more accuracy and robustness in protein cell abundance quantification compared to TCA two-step approach, regardless of the step size used in PGD optimization. Our algorithm also identifies the proteins contributing to the latent cellular heterogeneity in bulk multi-omics profiles, which may elucidate the cell marker proteins for certain tissue types. This multimodal deconvolution framework is more favorable in population-scale studies compared to the single modal deconvolution or single cell profiling, since it leverages distinct molecular sources and does not require unsupervised cell clustering or discrimination.

Another potential utility of our tool is the high-resolution purification of individualized pure cell expression, which is an essential component in the output of our algorithm and paves the way for deep proteome profiling at single cell type resolution. However, the current implementation of joint deconvolution algorithm emphasizes the accuracy of cell abundance quantification, which may sacrifice the power of high-resolution purification. A possible solution to precisely recover individualized multimodal pure cell expression is to train a deep learning model (such as autoencoder [21]) on the observed bulk proteomes and the reference panels pre-estimated by MICSQTL algorithm. Hence, it’s worth employing Stochastic Gradient Descent in the future to reduce the errors in pre-estimated reference panels.

Altogether, the joint deconvolution algorithm substantially improves the cell abundance estimation in bulk proteome by integrating modalities and using PGD for high-dimensional parameter optimization. The impact of PGD optimization on the quantified cell abundances and individualized purification will be assessed extensively in future. Our tool MICSQTL not only fills the methodological gap in bulk proteomics deconvolution and multimodal data integration, but also sheds light on the design of a comprehensive experiment that profiles single cell MS proteomes matched to the bulk multimodal samples.

## Methods

### Multimodal joint deconvolution

For the tissue biospecimen of individual *i*, we measure the expressions of protein *g* (*g* = 1, …, *G*) and mRNA transcript (or gene) *m* (*m* = 1, …, *M*), respectively, denoted by 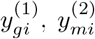. The unobserved high-resolution pure cell reference panels are denoted by 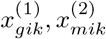, individually. The molecular source-specific cellular fractions for cell type *k* are 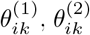 (*k* = 1, …, *K*), determined by the common tissue-specific cell counts (fractions) *p*_*ik*_ and source-specific cell size factors 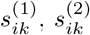. That is 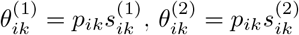. Thus, the bulk multi-modal expression data are modeled as

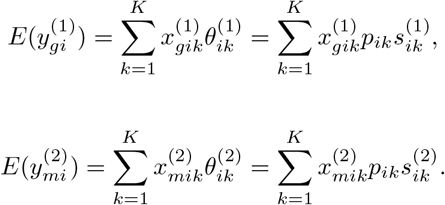

We denote 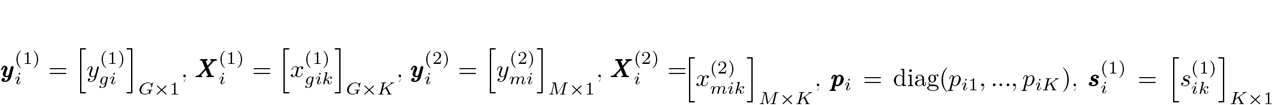 and 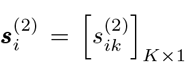 .The above source-specific models are linked by the common tissue-specific cell counts fractions ***p***_*i*_. Hence, we propose to jointly estimate the high-dimensional non-negative parameters 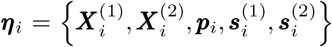 by minimizing a loss function ***L***(***η***_*i*_) that integrates 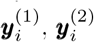. This is achieved by solving

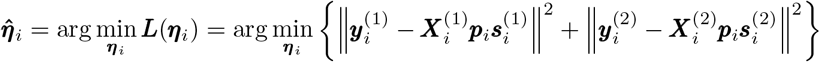

subject to min*{****η***_*i*_*} ≥* 0.

This loss function joins individual linear models by the shared cell counts ***p***_*i*_ as an extension of the Non-negative Matrix Factorization. The sample-wise high-dimensional non-negative parameters are estimated by Projected Gradient Descent [22] optimization:

1. Initialize 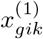 and 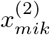 by sampling from Normal distributions *N* (*μ*^(1)^, *σ*^(1)^) and *N* (*μ*^(2)^, *σ*^(2)^), where *μ*^(1)^ and *σ*^(1)^, *μ*^(2)^ and *σ*^(2)^ are mean and standard deviation derived from 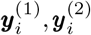. Initialize ***p***_*i*_ by Dirichlet distribution *Dir*(***α***_0_), where ***α***_0_ is derived from an external reference cell counts. Set 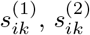 initials at ones.
2. Evaluate the loss function’s first gradient at current estimates: 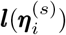.
3. Update parameters with non-negative bounds: 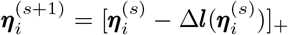 where Δ is step size.
4. Repeat steps 2-3 until convergence: max {|***η***^(*s*+1)^ − ***η***^(*s*)^|} *< ϵ* or a limit of iterations (e.g., 500).
5. Normalize the cellular fractions as 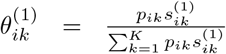 and 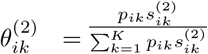

### Integrative signature selection

The decomposition provided by AJIVE enables the identification of underlying biological patterns that are common to all molecular modalities. Suppose there are *J* molecule sources for the same sets of *N* samples but different features from each source. For the bulk expression matrices ***Y*** ^(*j*)^ (*j* = 1, …, *J*) in distinct sources, AJIVE integrates ***Y*** ^(*j*)^’s to reconstruct each by three components: ***Y*** ^(*j*)^ = ***C***^(*j*)^ + ***I***^(*j*)^ + ***E***^(*j*)^, where ***C***^(*j*)^ represents the common variation originating from the *j*th modality, ***I***^(*j*)^ and ***E***^(*j*)^ are the source-specific structured variation and the residual noise, respectively.

The top proteins contributing to the common variation shared between proteome (***Y*** ^(1)^) and signature genes (***Y*** ^(2)^) are selected by using the loadings ***V*** ^(1)^ obtained from the singular value decomposition (SVD) of matrix ***C***^(1)^ = ***V*** ^(1)^***D***^(1)^***U*** ^(1)^, where ***V*** ^(1)^ is *G× r* with *r* as reduced rank estimated by the Wedin bound procedure [12]. Next, we compute 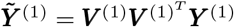 as a projected approximation to the observed bulk proteome with rank *r*. For each protein *g*, we calculate the distance between 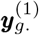and 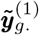 by the Euclidean distance 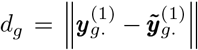. To select the proteins that contribute most significantly to the shared variation, we choose the proteins with the smallest distances *d*_*g*_.

### Cell-type-specific protein QTL mapping

The potential errors produced in sample-wise deconvoluted proteomes may lead to bias in cell-type-specific QTL mapping. Hence, we apply the method implemented in TOAST [23] to perform cell-type-specific protein differential analysis for genotypes based on the bulk proteomes, estimated protein cellular composition, and whole genome sequencing variants. This method performs F-test on the hypothesis for the cell-type-specific changes across three genetic groups [24], with false discovery rate being well controlled according to the previous simulation studies [25, 26].

## Declarations

### Data availability

The CITE-seq data is available on GEO under accession number GSE164378. The mass spectrometry data and RNA-seq data used in this study are available in the Synapse database under accession code syn26231732.

### Code availability

- MICSQTL is a Bioconductor package with license GPL-3, available at https://bioconductor.org/packages/MICSQTL.
- CIBERSORT (https://cibersortx.stanford.edu/)
- TOAST (https://bioconductor.org/packages/TOAST)
- TCA (https://cran.r-project.org/web/packages/TCA)

### Authors’ contributions

Y.P., X.W., J.P., and Q.L. conceive this study. Y.P. develops the algorithms and R package as maintainer, performs simulation and real data analysis, and visualizes analysis results. Q.L. proposes the algorithm and designs R package, simulation study, and real data analysis. Y.P. and Q.L. write the manuscript. X.W., C.L., and J.P. generate and share the real data, contribute to methodology discussion, and interpret real data analysis results. All authors reviewed the manuscript.

