## Supplementary File for "Multimodal joint deconvolution and integrative signature selection in proteomics"

Yue Pan, Xusheng Wang, Chunyu Liu, Junmin Peng, and Qian Li*

### Simulation design for tissue-matched transcriptome-proteome

To benchmark our algorithm, we synthesize matched bulk transcriptome-proteome with ground truth of cellular fractions of astrocytes, exitatory neurons, microglia, and oligodendrocytes in each molecular source, using parameters estimated from a public scRNA-seq data [1]. We first generate the transcript proportions $\boldsymbol{\theta}_i^{(1)}$ from the Dirichlet distribution with parameters for control group ($\bar{\boldsymbol{\alpha}}_C$) and diseased group ($\bar{\boldsymbol{\alpha}}_D$), respectively. The parameters $\bar{\boldsymbol{\alpha}}_C$ and $\bar{\boldsymbol{\alpha}}_D$ are estimated from real human prefrontal cortex snRNA-seq data [1], while the protein proportions $\boldsymbol{\theta}_i^{(2)}$ is obtained by random shifting $\theta_{ik}^{(2)} = \theta_{ik}^{(1)} c_{ik}$, whereas $c_{ik} \sim \text{Unif}(0.9, 1.1)$ with constraint $\sum_{k=1}^{K} \theta_{ik}^{(2)} = 1$. The correlation coefficient between synthesized transcript and protein proportions varies across cell types.

The expected mRNA expression and overdispersion for subject $i$ and cell type $k$ are simulated by the multivariate normal (MVN) distribution $\boldsymbol{x}_{ik}^{(1)} \sim MVN(\tilde{\boldsymbol{\mu}}_k, \tilde{\boldsymbol{\Sigma}}_k)$, $\boldsymbol{\Phi}_{ik} \sim MVN(\bar{\boldsymbol{\mu}}_k, \bar{\boldsymbol{\Sigma}}_k)$, where $\tilde{\boldsymbol{\mu}}_k, \bar{\boldsymbol{\mu}}_k, \tilde{\boldsymbol{\Sigma}}_k, \bar{\boldsymbol{\Sigma}}_k$ are estimated from the same snRNA-seq data. For each cell type $k$, the true reference mRNA expression in subject $i$ is generated by Gamma distribution $\lambda_{gik} \sim \Gamma(\exp(-\Phi_{gik}), x_{gik}^{(1)} \exp(\Phi_{gik}))$. The true mixture-cell mRNA expression level per sample is $\bar{\lambda}_{gi} = \boldsymbol{\lambda}'_{gi}\boldsymbol{\theta}_i^{(1)}$, where $\boldsymbol{\lambda}'_{gi} = (\lambda_{gi1}, ..., \lambda_{giK})$ and $\bar{\lambda}_{gi}$ still follows Gamma distribution [2]. The observed bulk RNA-seq counts is sampled from the Poisson distribution $\boldsymbol{y}_{gi}^{(1)} \sim Pois(\bar{\lambda}_{gi})$.

To generate pure cell proteomics data, the purified proteomes from human brain tissues are needed but lacking. Hence, we used a public pure cell mouse brain proteomes [3] as surrogate reference to acquire the mean expression $\boldsymbol{\mu}_k$ for each protein and the Protein-Protein Interaction structure (variance-covariance matrix) $\boldsymbol{\Sigma}_k$ within each cell type $k$. Note that the cell types used in aforementioned mRNA expression simulation are available in the mouse brain purified proteomes. Then for individual $i$, the cell-type-specific protein expression is simulated by $\boldsymbol{x}_{ik}^{(2)} \sim MNV(\boldsymbol{\mu}_k, \boldsymbol{\Sigma}_k)$. The cell-type-specific reference protein expression $x_{gik}^{(2)}$ is generated by truncated Normal distribution to mimic the limit of detection (LOD) in mass spectrometry, i.e.,

$$z_{gi} \sim \max\{\text{LOD}, \boldsymbol{x}_{gi}^{(2)}\boldsymbol{\theta}_i^{(2)}\}$$

The mixture-cell protein expression is sampled from $\boldsymbol{y}_{gi}^{(2)} \sim N(z_{gi}, \sigma_0^2)$, where $\sigma_0^2$ is estimated from the real bulk proteomics data.

### Human brain tissue

A total of $N = 264$ postmortem frozen human frontal cortex tissue samples from the Stanley Medical Research Institute (SMRI) and Banner Sun Health Research Institute (BSHRI) were used for this study. These samples were collected from $N = 194$ neurotypical controls, $N = 45$ individuals with schizophrenia (SCZ), and $N = 25$ individuals with bipolar disorder (BP). The tissues were weighed and homogenized in lysis buffer with a 1×PhosSTOP phosphatase inhibitor cocktail (Sigma-Aldrich). The total protein concentration of each sample was measured by the BCA Protein Assay Kit (Thermo Fisher Scientific), and confirmed by Coomassie-stained short SDS gels.

### Identification and quantification by JUMP software suite

We performed peptide identification with the JUMP search engine [4] to improve the sensitivity and specificity. JUMP searched MS/MS raw data against a composite target/decoy database to evaluate FDR. The target human protein sequences (83,955 entries) were downloaded from the UniProt database. The decoy database was generated by reversing to generate a decoy database that was concatenated to the target database. FDR was estimated by the ratio of the number of decoy matches and the number of target matches. Putative PSMs were filtered by mass accuracy and then grouped by precursor ion charge state and filtered by JUMP-based matching scores to reduce FDR below 1% for proteins during the whole proteome analysis. If one peptide could be generated from multiple homologous proteins, based on the rule of parsimony, the peptide was assigned to the canonical protein form in the manually curated Swiss-Prot database. We first extracted TMT reporter ion intensities of each PSM and corrected the raw intensities based on isotopic distribution of each labeling reagent. We discarded PSMs with low intensities (i.e., the minimum intensity of 1,000 and median intensity of 5,000). After normalizing abundance with the trimmed median intensity of all PSMs, we calculated the mean-centered intensities across samples (e.g., relative intensities between each sample and the mean) and summarized protein relative intensities by averaging related PSMs. Finally, we derived protein absolute intensities by multiplying the relative intensities by the grand mean of the three most highly abundant PSMs.

### RNA-seq transcriptomics preprocessing

We used different RNA preparation techniques for human brain samples from SMRI and BSHRI. For SMRI brain samples, total RNA was isolated for SMRI samples through organic extraction. Total RNA was precipitated with isopropanol at room temperature, pelleted, washed with 75% ethanol, and resuspended in DEPC treated water. For BSHRI brain samples, total RNA was mixed with ethanol and applied to a miRNeasy mini-column. Columns were treated with the RNase-free DNase digestion set (Qiagen), then washed with the appropriate miRNeasy mini kit buffers. All total RNA samples that passed QC for library generation were assayed by the Qubit 2.0 RNA BR Assay or Bioanalyzer RNA 6000 Nano assay kit. All FASTQ files were trimmed for adapter sequence and low base call quality (Phred score $< 30$ at ends) using cutadapt (v1.12) and then aligned to the GRCH37 (i.e., hg19) reference genome with STAR (2.4.2a) using GENCODE gene annotations. BAM files were sorted using samtools (v1.3). Gene expression levels were quantified using RSEM (v1.2.29).

### Additional results in simulation and real data application

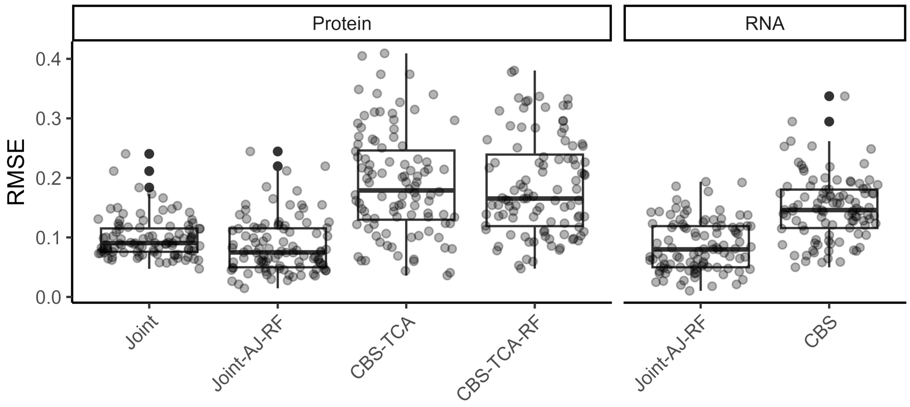


Figure S1: Simulation scenario A with PGD optimization step size $\Delta = 10^{-6}$ for cell size factors in joint deconvolution and inaccurate signature matrix in CBS and TCA.

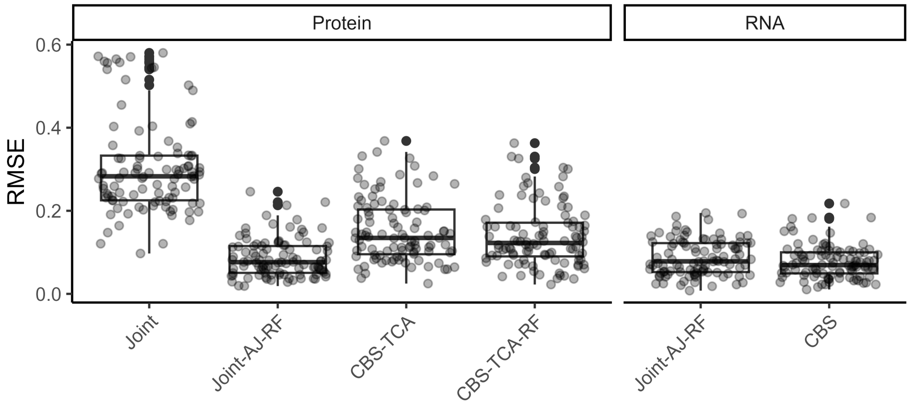


Figure S2: Simulation scenario A with PGD optimization step size $\Delta = 10^{-5}$ for cell size factors in joint deconvolution and accurate signature matrix in CBS and TCA.

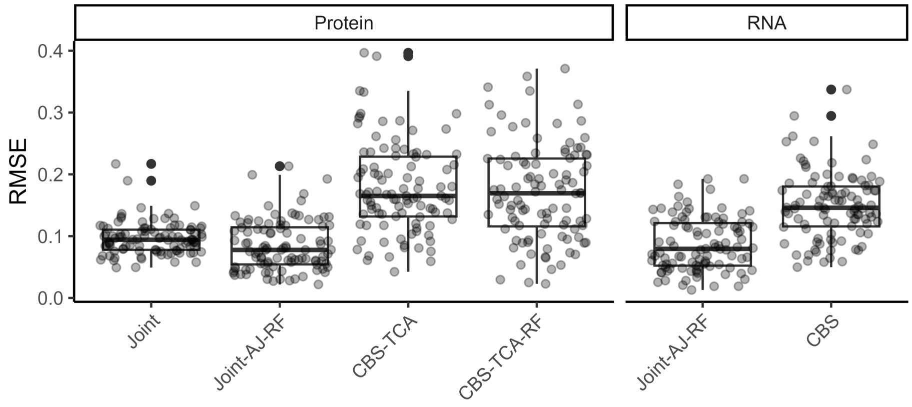


Figure S3: Simulation scenario B with PGD optimization step size $\Delta = 10^{-6}$ for cell size factors in joint deconvolution and inaccurate signature matrix in CBS and TCA.

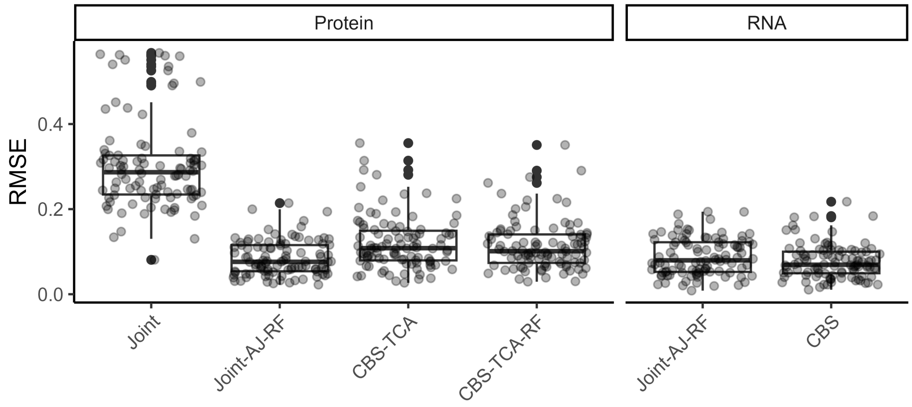


Figure S4: Simulation scenario B with PGD optimization step size $\Delta = 10^{-5}$ for cell size factors in joint deconvolution and accurate signature matrix in CBS and TCA.

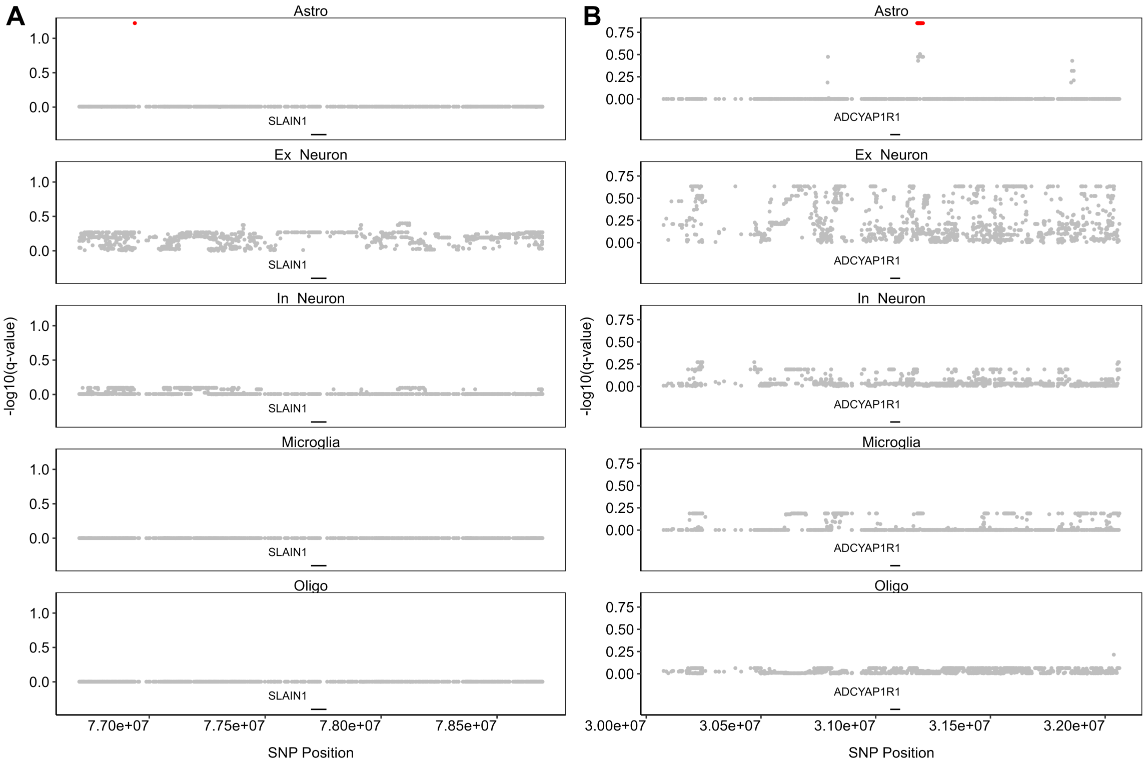


Figure S5: FDR-controlled q-value for cell-type-specific protein quantitative trait loci (pQTL) analysis for proteins (A) SLAIN1 and (B) ADCYAP1R1.
